# Skeletal muscle tissue secretes more extracellular vesicles than white adipose tissue and myofibers are a major source ex vivo but not in vivo

**DOI:** 10.1101/2020.09.27.313932

**Authors:** Andrea L. Estrada, Zackary Valenti, Gabriella Hehn, Christopher P. Allen, Nicole A. Kruh-Garcia, Daniel S. Lark

**Author notes:** Corresponding Author: Daniel Lark, Ph.D.

## Abstract

Circulating extracellular vesicles (EVs) are biomarkers of and contributors to the etiology of disease. Skeletal muscle (SkM) and white adipose tissue (WAT) are the two largest organs by mass in humans and rodents but the relative contribution of EVs from these tissues is unknown. We hypothesized that SkM tissue secretes more EVs than WAT and that a dual fluorescent reporter mouse could be used to detect SkM myofiber-derived EVs *in vivo*. Human Protein Atlas data and directly measuring EV secretion in mouse SkM and WAT using an *ex vivo* tissue explant model confirmed that SkM tissue secretes more EVs than WAT. Differences in EV secretion between SkM and WAT were not due to SkM contraction but may be explained by differences in tissue metabolic capacity. A SkM myofiber-specific dual fluorescent reporter mouse was created. Spectral flow cytometry revealed that SkM myofibers are a major source of SkM tissue-derived EVs *ex vivo* but few reach the circulation *in vivo*. Our findings demonstrate that SkM secretes more EVs than WAT and many come from SkM myofibers, but our *in vivo* data indicate that EVs secreted by SkM myofibers may remain primarily in their local extracellular environment.

## Background and Rationale

Extracellular vesicles (EVs) are a broad class of cell-secreted particles found in biofluids throughout the body. Circulating EVs are both biomarkers and contributors to human metabolic disease etiology (26). Skeletal muscle (SkM) and white adipose tissue (WAT) are the two largest organs (by mass) in the body and are central to the etiology of diseases like obesity and Type 2 diabetes. Their mass and disease relevance have led to studies investigating whether their EVs play a causal role in the development of insulin resistance, a component of the metabolic syndrome. High fat feeding, a common method for causing insulin resistance in rodents and humans, causes an increase in EV secretion from cultured SkM tissue and WAT explants (1, 7). Furthermore, injecting EVs from SkM tissue or WAT explants of diet-induced obese mice into lean recipient mice promote insulin resistance (1, 5). These data suggest that exogenous SkM and WAT-derived EVs play a role in metabolic disease etiology. However, to date there have been no direct comparisons of EV secretion by SkM or WAT tissues nor studies exploring to what extent SkM tissue-derived EVs originate from SkM myofibers.

Very little data exists regarding whether, and to what extent, non-vascular cells are able to secrete EVs that reach the circulation. Most of this emerging knowledge comes from transgenic fluorescent reporter mice. For example, the “mT/mG” (referred to as mG/mT mice in the current report) conditional dual fluorescent reporter mouse has been used to demonstrate that bronchoalveolar epithelial cells secrete EVs that can be found in the lung but not serum (28). Studies using WAT transplantation suggest that adipocyte-derived EVs can transport micro RNAs (miRNAs) to distal organs and tissues (32) while studies using mice with adipocyte-specific expression of tdTomato suggests that adipocytes are not a major contributor to circulating EVs (7). To our knowledge, there are no published studies demonstrating the existence of SkM myofiber-derived EVs in blood circulation *in vivo*. Furthermore, the biophysical mechanism whereby EVs secreted from non-vascular cells travel from the local extracellular environment across the vascular endothelium into the blood has yet to be described.

EVs from SkM cells are principally understood based on monocultures of proliferating myocytes and, to a lesser extent, differentiated myotubes (30). Proliferating C2C12 myocytes secrete EVs that improve endothelial function *in vitro* (25). EV secretion from C2C12 myocytes is increased by the saturated fatty acid palmitate and these palmitate-induced C2C12 myocyte-derived EVs impair insulin signaling in recipient cells (18). While these studies have improved our understanding of EV biology in SkM cells, proliferating myocytes do not necessarily behave like the terminally differentiated myofibers that dominate SkM tissues. Proliferating myocytes are primarily glycolytic whereas SkM myofibers in tissues have a metabolic phenotype determined in large part by fiber type (i.e. fast versus slow twitch). It is therefore unsurprising that EVs from proliferating SkM myocytes have a distinct composition compared to EVs secreted by differentiated SkM myotubes (8). These data demonstrate a putative role for SkM-derived EVs in intercellular and interorgan communication and suggest that oxidative metabolism may be a determinant of EV secretion.

Cell monocultures are a relatively simple system for studying SkM myocyte-derived EVs, but they lack the organization and extracellular matrix characteristic of tissues. *Ex vivo* incubation of SkM tissue explants has been shown to be an alternative approach to study EVs (1, 4, 18, 25). This method yields EVs with a comparable size, morphology and composition to EVs from cultured cells or blood (1). SkM tissue-derived EVs from obese mice increase expression of genes involved in regulating beta-cell mass in lean recipient mice (18). SkM tissue-derived EVs from hind limb denervated mice alter adipocyte gene expression in a co-culture system (4). In addition to SkM explants, WAT explants have also been studied. EVs expressing a white adipocyte profile for adipokines and adipocyte-derived proteins have been found in human plasma (3). In addition, the WAT explant model has been employed to characterize adipocyte-specific EV-like particles (5). These findings demonstrate the utility of tissue explants to measure EV secretion in specific tissues and to compare EV secretion between tissues.

In this report, we examined curated human protein and gene expression data and performed *ex vivo* measurements of EV secretion to test the hypothesis that SkM tissue secretes more EVs than WAT per unit of mass. We then generated a SkM myofiber-specific mG/mT mouse to test the hypothesis that SkM myofibers within SkM tissue secrete EVs *ex vivo* and *in vivo*.

## Methods

### Secondary Analysis of Data from the Human Protein Atlas

A retrospective analysis of publicly available protein and gene expression data from the Human Protein Atlas version 19.3 (HPA) was completed. Comparisons were made using immunohistochemistry (skeletal muscle myocytes versus white adipose tissue adipocytes) and RNA-Seq (skeletal muscle versus adipose tissues) from the “Normal Tissue Data” and “RNA consensus tissue gene data” files, respectively. Specifically, we compared the twenty-three genes annotated in the “Extracellular Vesicle Biogenesis” gene ontology cellular process (GO:0140112). IHC data is reported by HPA in four ordinal variables that were assigned numbers (Not Detected = 0, Low = 1, Medium = 2 and High Expression = 3). RNA data is reported as normalized expression, aggregating RNA-seq data processed using three different analytical workflows (HPA, GTEx and FANTOM). All data for all genes are provided in **Table 1**. Because these data are not reported as replicates, no statistical comparisons were possible. Instead, ordinal IHC data were either greater in SkM, greater in WAT, or comparable. Continuous gene expression data were either greater in SkM or greater in WAT.

**Table 1:** SkM is enriched in EV Biogenesis proteins compared to WAT based on IHC and RNA-Seq data from The Human Protein Atlas. Data obtained from **The Human Protein Atlas** Version 19.3 (www.humanproteinatlas.org). IHC expression values were obtained from the “normal tissue data” file. Data were converted from text to ranked values (Not Detected = 0, Low = 1, Medium – 2 and High Expression = 3). RNA-Seq data was obtained from the “RNA consensus tissue gene data” file. These data are reported as normalized expression aggregated across three distinct RNA-Seq workflows (HPA, GTEx and FANTOM). The specific genes analyzed comprise the gene ontology cellular process “EV Biogenesis” (GO:0140112).

| Protein/Gene | IHC in Cell Types |  |  | RNA-Seq in Tissues |  |  | IHC + RNA |
| --- | --- | --- | --- | --- | --- | --- | --- |
|  | Adipose Tissue | SkM | Greater | White Adipose Tissue | SkM Tissue | Greater |  |
| Arrdc1 | 0 | 3 | SkM | 8.4 | 9.1 | SkM | Yes |
| Arrdc4 | 1 | 3 | SkM | 9.2 | 10.1 | SkM | Yes |
| Atp13a2 | 0 | 2 | SkM | 9.6 | 7.5 | Adipose | No |
| Cd34 | 0 | 0 | Equal | 61.9 | 39.7 | Adipose | No |
| Chmp2a | 1 | 1 | Equal | 40.7 | 75.7 | SkM | No |
| Cops5 | 3 | 3 | Equal | 20.2 | 64 | SkM | No |
| Hgs | 0 | 2 | SkM | 26.8 | 36.3 | SkM | Yes |
| Myo5b | 0 | 0 | Equal | 2 | 0.6 | Adipose | No |
| Pdcd6ip | 0 | 2 | SkM | 39.1 | 29.5 | Adipose | No |
| Prkn | 0 | 2 | SkM | 6.8 | 41.8 | SkM | Yes |
| Rab7a | 2 | 2 | Equal | 61.5 | 146.3 | SkM | No |
| Rab11a | 0 | 1 | SkM | 43.3 | 27.3 | Adipose | No |
| Rab27a | 0 | 0 | Equal | 11.4 | 5.1 | Adipose | No |
| Sdc1 | 0 | 0 | Equal | 0.8 | 0.5 | Adipose | No |
| Sdc4 | 2 | 2 | Equal | 15.9 | 25.5 | SkM | No |
| Sdcbp | 0 | 0 | Equal | 106.2 | 22.5 | Adipose | No |
| Smpd3 | 0 | 0 | Equal | 3.5 | 1.8 | Adipose | No |
| Snf8 | 0 | 1 | SkM | 22.5 | 40.7 | SkM | Yes |
| Stam | 1 | 0 | Adipose | 13.5 | 26 | SkM | No |
| Steap3 | 2 | 1 | Adipose | 8.1 | 10.9 | SkM | No |
| Tsg101 | 0 | 2 | SkM | 24.7 | 44.5 | SkM | Yes |
| Vps4a | 1 | 1 | Equal | 20.2 | 81.6 | SkM | No |
| Vps4b | 0 | 0 | Equal | 16.8 | 9.9 | Adipose | No |
|  |  | SkM | 9 |  | SkM | 13 | Targets |
|  |  | Equal | 12 |  | WAT | 10 | 6 |
|  |  | WAT | 0 |  |  |  |  |

### Mouse Models and Studies

All studies were approved by the Colorado State University Institutional Animal Care and Use Committee (Protocol #: 18-7858A). All mice were on the C57BL/6J background and originally purchased from Jackson Laboratories. Wild type mice were allowed a 2-week acclimation period prior to commencing studies. A colony of “mT/mG” (referred to as mG/mT mice in this report) fluorescent reporter mice was established using (B6.129(Cg)-Gt(ROSA)26Sortm4(ACTB-tdTomato,-EGFP)Luo/J – Stock #: 007676) mice. These mice express membrane-targeted tandem dimer Tomato (mT) in all cells in the absence of Cre recombinase. In cells expressing Cre recombinase, the mT gene construct is excised so a membrane-targeted enhanced green fluorescent protein (mG) (24) is expressed instead. For our experiments, mice with expression of SkM myocyte-specific expression of mG were desired. This was achieved by crossing mG/mT mice with HSA-Cre79 (B6.Cg-Tg(ACTA1-cre)79Jme/J – Stock #: 006149) mice and the progeny of those mice (+/− Cre, +/− mG/mT) bred back to homozygous (+/+) mG/mT mice. Tail snip or ear punch DNA was analyzed via PCR to verify the mG/mT and Cre alleles. Male mice with homozygous expression of the “mG/mT” allele were bred to female mice homozygous for the mG/mT allele and heterozygous for HSA-Cre. Progeny were either Cre negative and termed “mT mice” or Cre positive and termed “SkM-mG/mT mice”.

For plasma and tissue collection, mice were anesthetized with isoflurane and whole blood was collected via cardiac puncture. Whole blood was transferred into EDTA-coated tubes and immediately centrifuged at 5,000 x g at 4°C for 10 minutes to remove cells and debris. Plasma was collected by centrifugation at 18,000 x g at 4°C for 30 minutes and stored at −80°C. Vastus medialis and epididymal adipose tissue were subsequently collected and stored in ice-cold PBS until dissection.

### Ex Vivo EV Secretion Assay

This method was adapted from (1) with our specific approach illustrated in (**Figure 1**). Vastus medialis SkM and epididymal WAT were dissected into ~5 mg pieces in ice-cold PBS. SkM and WAT from male WT, mT and SkM-mG/mT mice (n=3 mice/genotype; ~20 weeks of age) was used for these experiments. ~50 mg of SkM tissue was placed in each well. Due to low EV yield from WAT, ~250 mg of WAT was used in each well. Tissue was cultured in poly-L-lysine coated 6 well plates (Corning Inc.) containing 2 mL of serum-free DMEM (Sigma D1145: 4.5 g/L D-Glucose and sodium bicarbonate, without L-Glutamine, sodium pyruvate or phenol red) and was supplemented with 1% penicillin/streptomycin. Cultured tissue was incubated for 24 hours in a 95% O_2_/ 5% CO_2_-supplemented incubator at 37°C. SkM and WAT tissues were collected from each well, weighed, snap frozen and stored at −80°C. EV-rich conditioned media was collected via centrifugation at 3,000 x g for 15 minutes, in order to clear cells and debris, and stored at −80°C until EV isolation.

**Figure 1:**
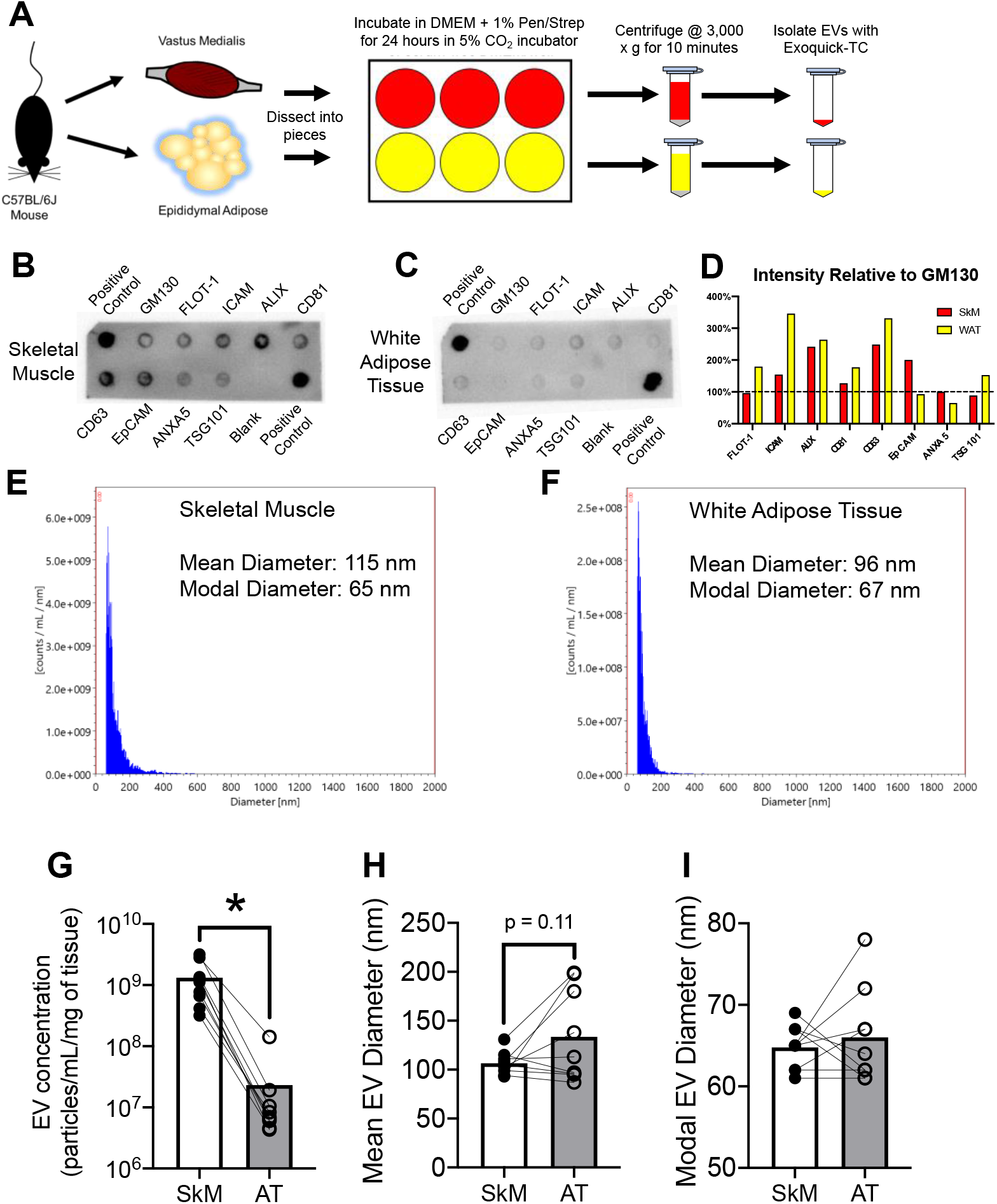
SkM secretes more EVs than WAT *ex vivo*. A) Assay workflow for measuring *ex vivo* EV secretion in mouse vastus medialis skeletal muscle (SkM) and epididymal white adipose tissue (WAT). Additional experimental details are provided in the ***Methods*** section. ExoCheck blots for EV-associated proteins in 25 ug of isolated B) SkM and C) WAT-derived EVs. D) Abundance of EV-associated proteins relative to GM130, a cytosolic protein not found in EVs. Representative NTA measurements from E) SkM and F) WAT-derived EVs. G-I) Comparison of EVs secreted by SkM versus WAT from male WT, mT and SkM-mG/mT mice over 24 hours. N=3 mice per genotype were pooled for analysis of n=9 total mice. G) Mass-adjusted EV secretion. H) Mean EV diameter. I) Modal EV diameter. * p < 0.01 using paired Student’s t-test.

Due to the smaller masses of soleus and plantaris compared to vastus, 12 wells plates with 0.75 mL of media per well was used to study these tissues. In all other respects, tissues were handled exactly the same as described above.

### Inhibition of Spontaneous SkM Contraction

To examine the role of spontaneous SkM contraction using the myosin ATPase inhibitor blebbistatin, 12 well plates with 0.75 mL of media were used. 10 μM blebbistatin or vehicle (DMSO) was added to each well to inhibit spontaneous SkM contraction as we have previously demonstrated (27). Tissues and media were subsequently handled in an identical fashion to the procedure described above.

### EV Isolation

EVs were isolated using Exoquick-TC or Exoquick Plasma according to the manufacturer’s instructions, or size exclusion chromatography as previously described (6). For *ex vivo* studies, conditioned media was combined with Exoquick-TC at a 5:1 ratio (i.e. 500 μl of media and 100 μl of Exoquick-TC) and incubated for 12 hours at 4°C. Samples were centrifuged at 1,500 x g for 30 minutes at 4°C and EVs were resuspended in 100 μL of 0.1 μM filtered PBS. NTA (described in detail below) was performed on freshly isolated EVs. Isolated EVs were stored at −80°C for subsequent analyses.

### Nanoparticle Tracking Analysis

Nanoparticle tracking analysis was performed on a three laser ViewSizer 3000 (Horiba Instruments). 0.1 μM filtered H_2_O was used as diluent and all EV concentrations were subtracted from a diluent file created specifically for samples run and analyzed that day. Focus was optimized for each individual sample immediately prior to recording. The following recording settings were used: Frame Rate = 30 fps, Exposure = 15 ms, Gain = 30 dB, Stir time between videos = 5 seconds, Wait time between videos = 3 seconds, Blue Laser Power = 210 mW, Green Laser Power = 12 mW, Red Laser Power = 8 mW, Frames per video = 300, Videos Recorded = 25, Temperature = 22°C. Videos were processed using ViewSizer software using the following settings as recommended by the manufacturer: Detection Threshold Type = Manual, Detection Threshold = 0.8, AutoThreshold = Disabled, Feature Radius = 40 pixels, Drift Correction = 0%. To avoid over or underestimation of particle concentration or size, samples were diluted in the same 0.1 μM filtered H_2_O as used to generate the diluent file in order to reach 120-150 tracks per frame as recommended by the manufacturer. EV concentration obtained from NTA was adjusted for dilution factor and then normalized to tissue mass. Therefore, EV concentration is reported as particles/mL/mg tissue.

### Protein Quantification

Protein from isolated EVs was analyzed using a micro BCA assay (Biorad) in polystyrene 96 well plates according to the manufacturer’s protocol on a SpectraMax M2 plate reader (Molecular Devices Inc.) at 562 nm absorbance.

### Exocheck Antibody Array

To determine the presence of EV-associated proteins (CD63, CD81, ALIX, FLOT1, ICAM1, EpCam, ANXA5 and TSG101) and cellular contamination (GM130) in EVs isolated from conditioned media, the Exocheck Antibody Array (Systems Bioscience) was used according to the manufacturer’s instructions. Due to low protein yield from WAT-derived EVs, 25 μg of protein measured using a micro BCA assay was used for both SkM and WAT-derived EVs. To image blots, the PVDF membranes were incubated for 2 minutes in SuperSignal West Pico Plus (Thermo Fisher Scientific) ECL reagent then imaged on a FluorChem E imager (Bio-Techne). Individual dots were traced with ImageJ and the intensity of each dot normalized to the intensity of the negative control dot.

### Flow Cytometry

Flow experiments were performed using a four laser (405, 488, 562 and 640 nm excitation) Aurora spectral flow cytometer (Cytek Biosciences) in the Colorado State University Flow Cytometry and Cell Sorting facility. EV detection and optimization was conducted in accordance with MISEV and MIFlowCyt guidelines (31, 34). Prior to each analysis, the cytometer was warmed up by running 0.2 μM-filtered water for 45 minutes. Quality control was performed using SpectroFlo^®^ Cytometer QC Beads (Cytek Biosciences, bead lot 2002) to normalize sensor gain as recommended by the manufacturer. Additionally, a clean flow cell procedure was conducted prior to sample analysis to minimize EV carryover. The flow cell was confirmed to be clean based on a low event rate (< 25-30 events/second) while running 0.1 μM filtered PBS. Side scatter was measured using the 405nm violet laser for improved detection of small EVs (21) and a side scatter threshold of 700 was applied unless otherwise noted. Finally, samples were run at a low flow rate (~15 μL/min), to minimize swarming and improve signal:noise. Additional optimization and validation experiments are described in the *<u>Results</u>* section. Data were collected in SpectroFlo^®^ and .fcs files were exported and further analyzed using FlowJo Version 10.6. To confirm that EV-associated fluorescence was from lipid vesicles, we compared isolated EVs analyzed as described above before and after detergent lysis via 0.1% NP-40 for 30 minutes at room temperature.

### Isolation of Mouse Splenocytes and Liver Cells

Splenocytes were isolated and purified as described on the R&D systems website (https://www.rndsystems.com/resources/protocols/leukocyte-preparation-protocol). Liver cells were isolated in an identical fashion but excluding the erythrocyte lysis step. Splenocytes and liver cells were re-suspended in PBS for flow cytometry.

### Isolation of Mouse Erythrocytes

Mouse erythrocytes were isolated and re-suspended as previously described (29). Whole blood was collected via cardiac puncture in heparinized tubes, centrifuged at 500 x g for 10 minutes at 4°C. Cells were then washed three times in “cell wash buffer” at 37°C (pH = 7.4) containing (in mM): 4.7 KCl, 2.0 CaCl_2_, 1.2 MgSO_4_, 140.5 NaCl, 21.0 Tris-base, 5.5 glucose and 0.5% BSA. Erythrocytes were re-suspended in bicarbonate-based buffer containing (in mM): 4.7 KCl, 2.0 CaCl_2_, 1.2 MgSO_4_, 140.5 NaCl, 11.1 D-Glucose, 23.8 NaHCO_3_ and 0.5% BSA for flow cytometry.

### Data Analysis

For all experiments, EV abundance and size are reported as mean +/− SEM. Statistical comparisons were performed using Prism 8 (GraphPad). Comparisons of EV abundance and diameter between tissues or in response to blebbistatin were made using paired Student’s t-test. Statistical comparisons were not possible for Human Protein Atlas data since replicate values are not provided.

## Results

### SkM is enriched in proteins involved in EV biogenesis compared to WAT in humans

We first tested the hypothesis that SkM has a greater capacity for EV secretion than WAT by performing a retrospective analysis of publicly available protein and gene expression data from the Human Protein Atlas (HPA) version 19.3. IHC and RNA-Seq data for the twenty-three genes from the “Extracellular Vesicle Biogenesis” gene ontology cellular process (GO:0140112) are provided in **Table 1**. Comparing IHC of SkM myofibers to WAT adipocytes revealed nine distinct proteins with greater relative expression in SkM myofibers. The remaining twelve proteins were comparable in expression; no proteins were greater in WAT adipocytes than SkM myofibers. RNA-Seq data revealed a total of thirteen genes more abundantly expressed in SkM; with the remaining 10 genes expressed to a greater extent in WAT than SkM. Combining results together revealed six EV biogenesis genes (Arrdc1, Arrdc4, Hgs, Parkin, Snf8 and TSG101) with greater protein and transcript in SkM compared to WAT. Overall, this secondary analysis of public data support the hypothesis that EV secretion is greater in SkM compared to WAT.

### SkM tissue secretes exponentially more EVs than WAT per unit of mass ex vivo

With evidence that SkM is enriched in EV biogenesis proteins relative to WAT, we directly compared EV secretion from SkM (vastus medialis) and WAT (epididymal visceral fat) of C57BL/6J mice using the *ex vivo* approach outlined in (**Figure 1A**). Dot blots of EVs isolated from SkM (**Figure 1B**) and WAT (**Figure 1C**) conditioned media show enrichment of EV-associated proteins (**Figure 1D**). Representative NTA analyses of EVs isolated from SkM (**Figure 1E**) and WAT (**Figure 1F**) conditioned media are provided and indicate a median diameter of ~70 nm consistent with small EVs.

We next compared EV abundance in the conditioned media from SkM tissue and WAT. EV abundance in conditioned media was on average one-hundred times greater in SkM compared to WAT after normalizing for tissue mass (**Figure 1G**). There was no difference in mean (**Figure 1H**) or modal EV diameter (**Figure 1I**) between SkM and WAT-derived EVs. These data show that SkM tissue, on a per mass basis, secretes more EVs than WAT in mice.

### Inhibiting spontaneous contraction does not decrease EV secretion from SkM explants

SkM is unique from most other tissues, including WAT, in its ability to contract. SkM contraction is thought to explain, at least in part, the increased abundance of circulating EVs observed during exercise (9, 35). SkM explants contract *ex vivo* but this can be prevented by the addition of 10 μM blebbistatin (27), a myosin ATPase inhibitor. We hypothesized that spontaneous contraction could explain the differences in EV secretion between SkM and WAT measured *ex vivo*. To test this hypothesis, we compared EV secretion from SkM explants in the presence of blebbistatin (10 μM) or an equal volume of DMSO vehicle. Contrary to our hypothesis, blebbistatin did not decrease SkM EV secretion (**Figure 2A**). However, blebbistatin did cause a decrease in mean (**Figure 2B**), but not modal (**Figure 2C**) EV diameter. These data indicate that greater EV secretion from SkM compared to WAT is independent of muscle contraction and instead possibly due to differences in metabolism.

**Figure 2:**
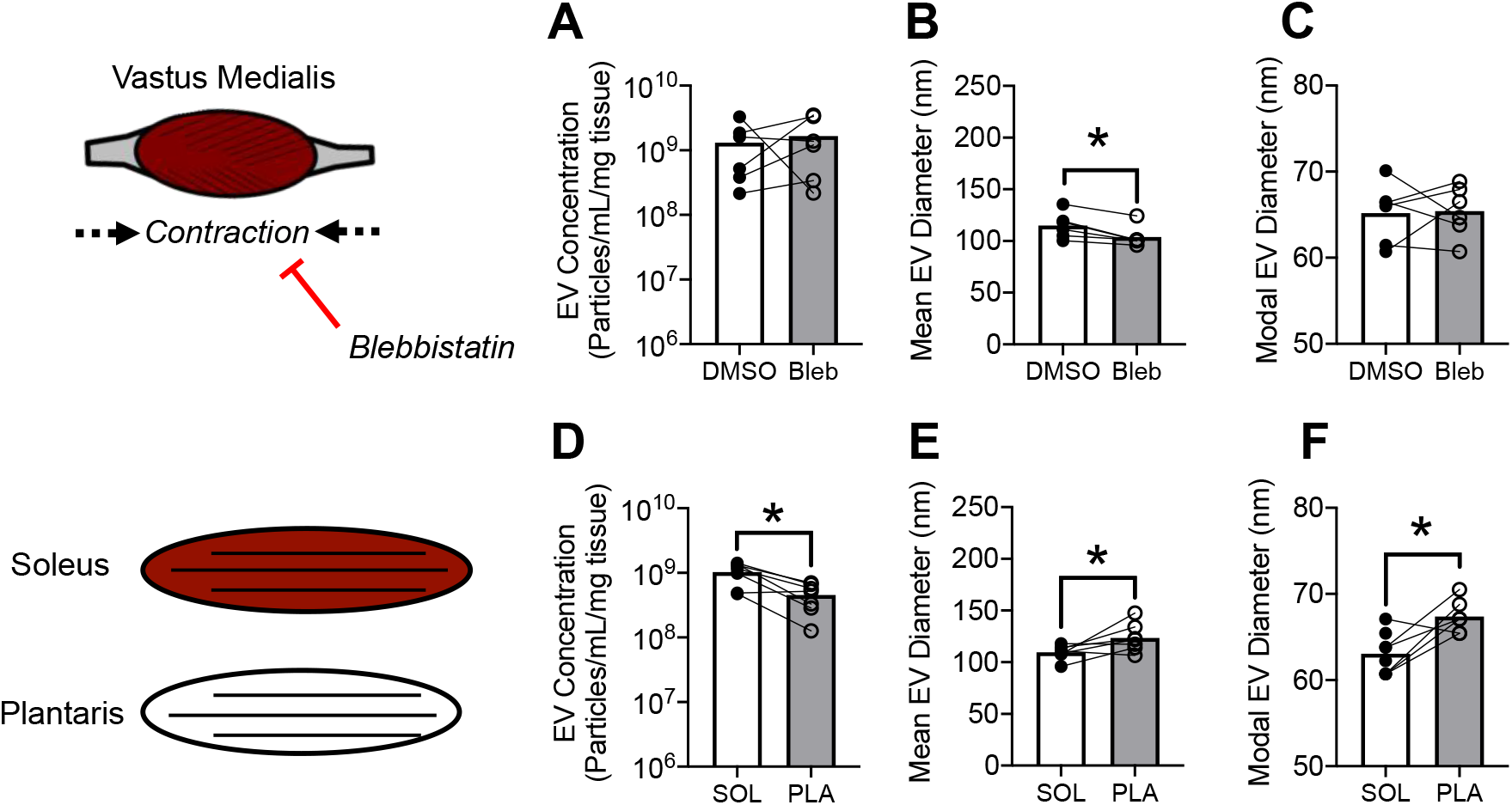
*Ex vivo* EV secretion is determined by metabolic capacity but not spontaneous contraction. A-C) Comparison of EVs secreted by vastus medialis skeletal muscle of male WT mice in the absence or presence of 10 μM blebbistatin or DMSO vehicle for 24 hours. A) Mass-adjusted EV secretion. B) Mean EV Diameter. C) Modal EV Diameter. N = 6 mice. * indicates p < 0.01 using paired student’s t-test. D-F) Comparison of EVs secreted by oxidative (soleus) or glycolytic (plantaris) skeletal muscle from female mT mice. D) Mass-adjusted EV secretion. E) Mean EV Diameter. F) Modal EV Diameter. N = 8 mice. * indicates p < 0.01 using paired student’s t-test.

### Metabolic capacity is a determinant of SkM EV secretion ex vivo

Since differences in EV secretion between SkM and WAT were not explained by contraction, we hypothesized that the intrinsic metabolic capacity of tissues might determine EV secretion. To test this hypothesis, we compared soleus, a highly oxidative muscle, to plantaris, a more glycolytic muscle. Consistent with our hypothesis, soleus secreted more EVs than plantaris per unit of mass (**Figure 2D**). Furthermore, soleus secreted EVs that were smaller both in mean (**Figure 2E**) and modal (**Figure 2F**) diameter compared to plantaris. These data suggest that metabolic capacity might explain differences in EV secretion between SkM and WAT.

### Spectral unmixing of enhanced GFP and tandem dimer Tomato

SkM tissue is a robust source of EVs, but it is not known to what extent specific cell types within SkM tissue secrete EVs. We developed a cell-specific dual fluorescent reporter mouse to determine the contribution of SkM myofibers to SkM tissue-derived EV secretion. In **Figure 3A**, we illustrate the genetic construct used to generate mice with ubiquitous expression of membrane-targeted tdTomato (Ex 554 nm/Em 581 nm) termed, “mT”. We then bred that line with mice expressing HSA-Cre, which have Cre recombinase expression specifically in SkM myofibers. The resulting mice have SkM myofiber-specific expression of membrane-targeted enhanced GFP (Ex 488 nm/Em 507 nm) termed, “mG”, and mT expression in all other cells. To detect mG and mT in individual particles, we utilized the spectral unmixing algorithm supplied in the SpectroFlo^®^ software to unmix eGFP, tdTomato and autofluorescence. Within this analysis, forty-eight detection channels collect data simultaneously and generate a spectral signature for each unique fluorophore. Spectral unmixing applies a modified least squares algorithm to not only extract autofluorescence, but also distinguish between fluorophores with and without overlapping spectra. Conventionally, *stained* and *unstained-control* cells are the required input to accomplish spectral unmixing. To unmix the spectra of tdTomato, splenocytes isolated from mT mice were used as the *stained* (fluorophore positive) population, and splenocytes from a WT mouse were used as an *unstained*-control (fluorophore negative) population. Monocytes were gated first on FSC-Area vs SSC-Area (**Figure 3B**) then on the presence of tdTomato (**Figure 3C**). The fluorescent spectral signature used for unmixing for tdTomato is shown in **Figure 3D**. To unmix eGFP, commercially available eGFP-labeled flow cytometer calibration beads (Takara Biosciences) and unstained beads (Thermo Fisher) were used. Beads were first gated based on FSC-Area vs SSC-Area (**Figure 3E**) then on the presence of eGFP (**Figure 3F**). The spectral signature for eGFP is shown in **Figure 3G**. All subsequent fluorescence data report tdTomato and eGFP (unmixed) signals.

**Figure 3:**
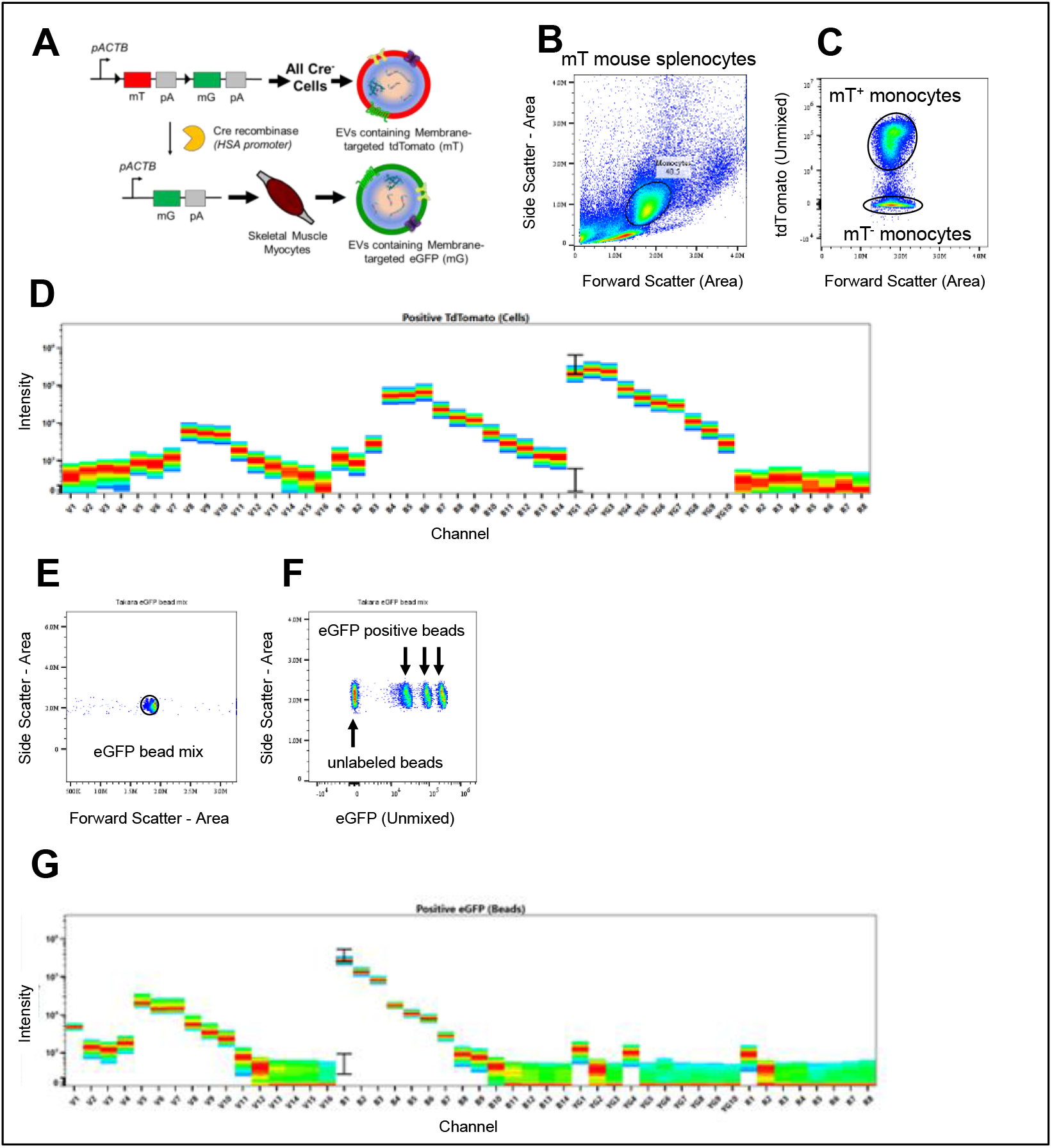
Spectral Unmixing of tdTomato and eGFP fluorophores. A) Gene editing approach predicting the secretion of tdTomato or eGFP-expressing EVs. B) Monocytes were isolated from mT mice using differential centrifugation and analyzed in a Cytek Aurora spectral flow cytometer. Monocytes were identified using SSC-Area vs FSC-Area. C) Monocyte populations were distinguished by gating on compensated tdTomato. D) Fluorescence spectra of tdTomato^+^ monocytes across 48 detection channels. E) eGFP-labeled beads were identified by plotting SSC-Area vs FSC-Area. F) eGFP+ and eGFP-beads were distinguished using compensated eGFP. G) Fluorescence spectra for eGFP^+^ beads across 48 detection channels.

### Detecting EV-sized particles using spectral flow cytometry

With the ability to discriminate between mT (tdTomato) and mG (eGFP) positive populations, the cellular origin of EVs derived from SkM tissue was determined. An optimized flow cytometry-based approach to detect EV populations based on side scatter using the 405 nm laser of a Cytek Aurora spectral flow cytometer was established. First, a side scatter threshold stepping analysis was performed using diluent 0.1 μM filtered PBS (**Figure 4A**). This analysis was performed to identify a threshold that eliminates the majority of instrument noise, thereby increasing the likelihood that detected events are biological particles like EVs.

**Figure 4:**
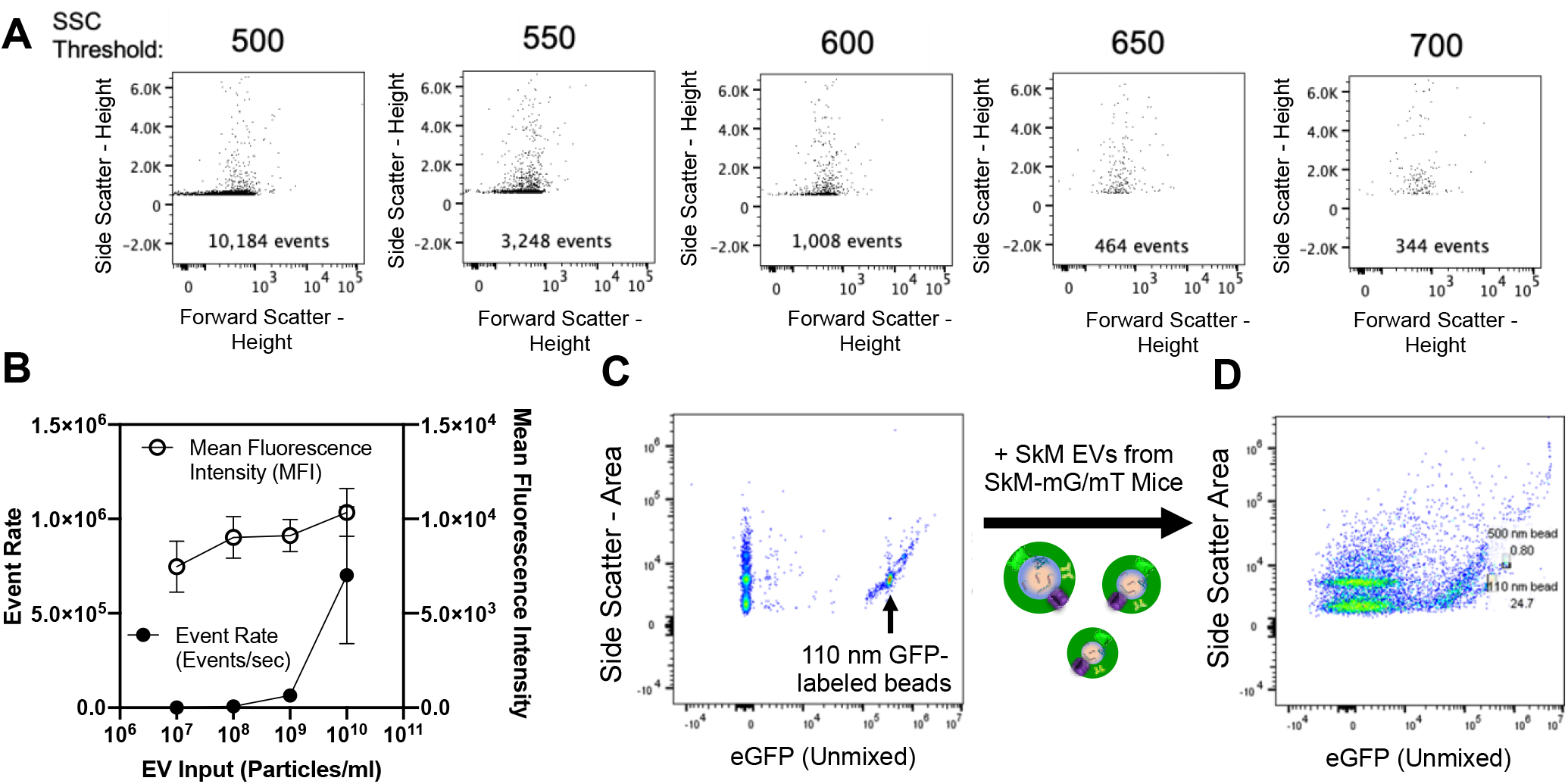
Detection of EV-sized particles using spectral flow cytometry. A) Side scatter threshold stepping using 0.1 μM filtered PBS was performed to minimize instrument noise. Inlaid units are events/second. B) Serial dilution of SkM-derived EVs from mT mice measuring mean fluorescence intensity (MFI) as a function of event rate (events/min). C) Detection of a mixture of Apogee beads containing both unlabeled silica and GFP-labeled polystyrene beads. D) Detection of Apogee beads spiked with SkM-derived EVs from SkM-mG/mT mice.

Next, co-incident detection (aka swarming) of particles was addressed. In order to increase the likelihood that mT^+^ or mG^+^ events represented single particles, serial dilutions of EVs were analyzed based on EV concentrations determined by NTA. Experiments began with 10^10^ EVs/mL. EVs were diluted an order of magnitude at each step. Swarming was considered minimal when the event rate decreased as a function of dilution while mean fluorescent intensity remained stable. **Figure 4B** summarizes a set of serial dilution experiments (n=3 mice) performed with SkM-derived EVs from SkM-mG/mT mice. Based on these results, it was determined that an EV concentration of 1×10^9^/mL or less was sufficient to minimize swarming. This concentration is consistent with previous flow studies using different flow cytometric instruments and particle compositions (11, 33).

Next we tested whether SkM-derived EVs from our fluorescent reporter mice could be detected using these established settings. To do this, we first confirmed the ability to detect a heterogenous mixture of small (110 nm diameter; indicated with arrow) GFP-labeled polystyrene beads (refractive index = 1.59) and similarly sized unlabeled silica beads (refractive index = 1.43) (**Figure 4C**). SkM-derived EVs from SkM-mG/mT mice were then added to the bead mixture and re-analyzed with the same settings. Shown in **Figure 4D**, SkM-derived EVs from SkM-mG/mT mice had similar side scatter properties to small GFP-labeled and unlabeled beads. Additionally, eGFP was detected across a range of side scatter intensities, suggesting the presence of fluorescently labeled EVs. Fluorescent EVs are further explored in **Figures 5 and 6**. These data demonstrate that both unlabeled and fluorescent EVs can be detected using spectral flow cytometry.

**Figure 5:**
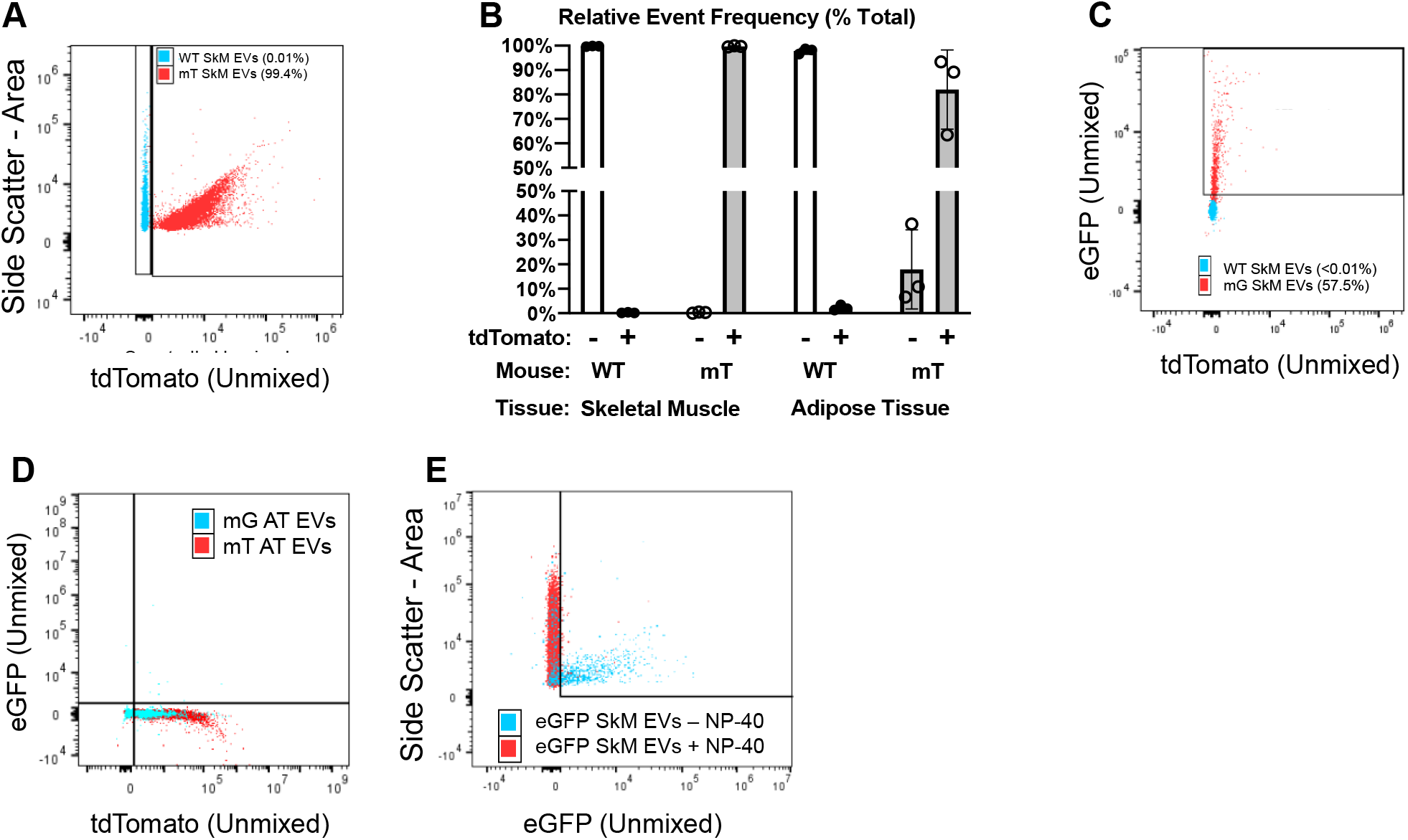
Detection of endogenously fluorescent EVs using spectral flow cytometry. A) Representative plot showing side scatter (y-axis) and tdTomato fluorescence (x-axis) in SkM EVs from WT (blue) and mT (red) mice measured at 10^8^ EVs/ml. B) Summary data for tdTomato^+^ events in EVs from SkM or WAT of WT or mT mice. C) Representative plot showing tdTomato (y-axis) and eGFP (x-axis) in SkM EVs from WT (blue) and SkM-mG/mT (red) mice. D) Representative plot showing tdTomato (y-axis) and eGFP (x-axis) in AT EVs from mT (blue) and SkM-mG/mT (red) mice showing no detectable presence of eGFP. E) Representative data showing that the lysing EVs with 0.5% NP-40 for 30 minutes at RT (red) causes the loss eGFP signal (x-axis) in SkM EVs from SkM-mG/mT mice. All data representative of three biological replicates per experiment.

**Figure 6:**
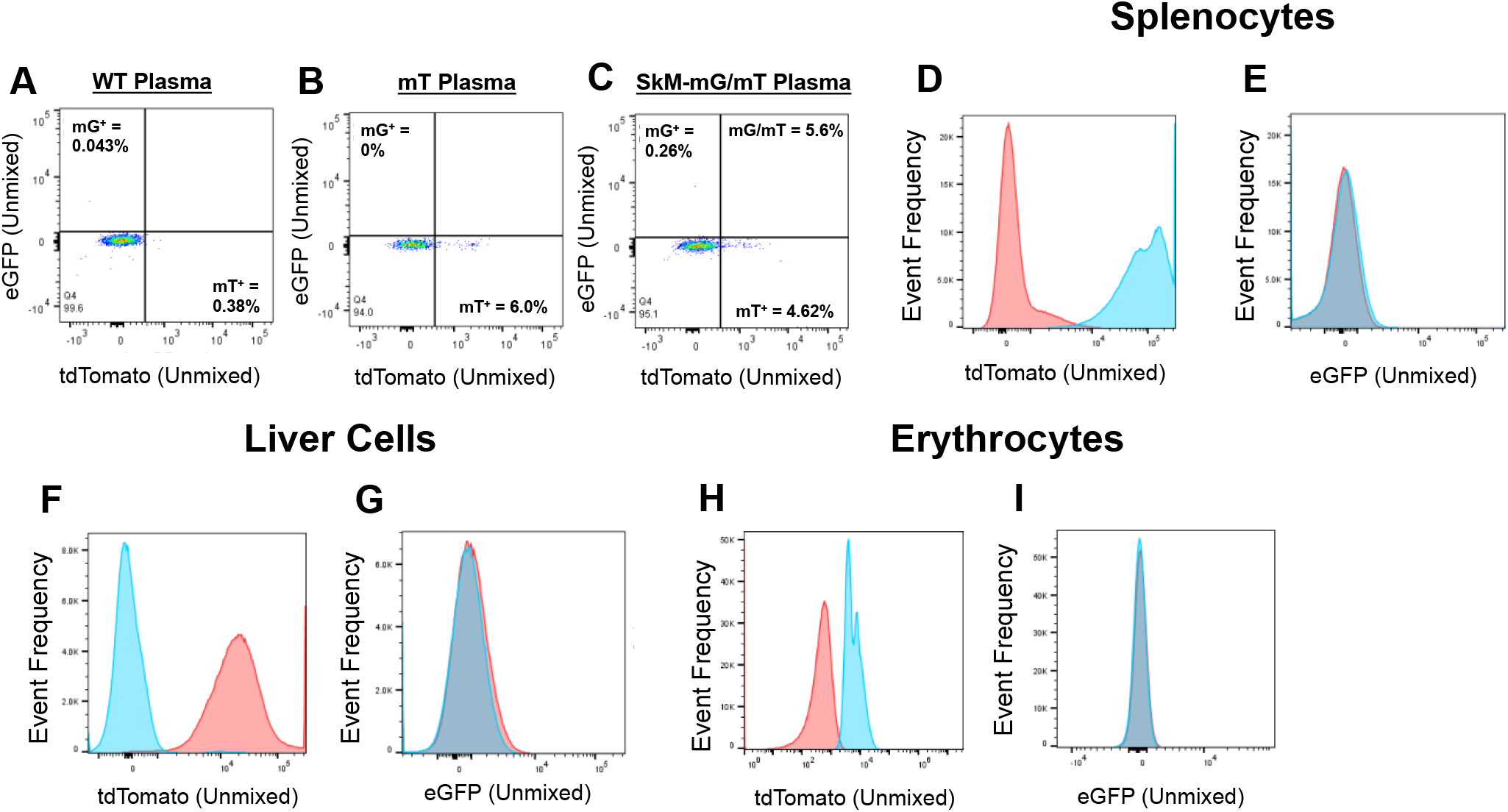
Evidence for systemic distribution of SkM myofiber-specific EVs *in vivo*. Plasma EVs from A) WT, B) mT and C) SkM-mG/mT mice were isolated using Exoquick-TC and analyzed at 1×10^11^ using spectral flow cytometry. Data are representative of EVs isolated with ExoQuick-TC (shown), ExoQuick Plasma or Size Exclusion Chromatography. D) Frequency histogram of tdTomato from splenocytes isolated from WT mice (red) or SkM-mG/mT mice (blue). E) Frequency histogram of eGFP from splenocytes isolated from WT (red) or SkM-mG/mT mice (blue). F) Frequency histogram of tdTomato in liver cells isolated from WT (blue) or SkM-mG/mT mice (red). G) Frequency histogram of eGFP in liver cells isolated from WT (blue) or SkM-mG/mT mice (red). H) Frequency histogram of tdTomato in erythrocytes isolated from WT (red) or SkM-mG/mT mice (blue). I) Frequency histogram of eGFP in erythrocytes isolated from WT (blue) or SkM-mG/mT mice (red).

### Endogenous fluorescent proteins are incorporated into SkM tissue-derived EVs secreted ex vivo

Equipped with the ability to detect EV-sized particles and independently resolve eGFP and tdTomato, we next measured the abundance of fluorescently-labeled EVs secreted by SkM at 10^9^ particles/mL. For these experiments, we used WT C57BL/6J mice (no fluorescence), mT mice (ubiquitous mT expression) or SkM-mG/mT mice (mG in SkM myocytes/myofibers, mT in all other cells). We first compared SkM EVs between WT and mT mice. As predicted, EVs isolated from WT SkM tissue lacked mT whereas nearly all EVs from mT mice (>98%) were mT^+^. An overlay of representative data from each genotype is provided in **Figure 5A**. Similar distributions were found in WAT analyzed at 10^7^ particles/mL due to the lower yield noted in **Figure 1G**. Summary data for these experiments is provided in **Figure 5B**. These results suggest that mT is incorporated into tissue-derived EVs secreted *ex vivo*.

Next, we tested the hypothesis that mG is incorporated into EVs in a SkM myofiber-specific fashion. On average, less than 0.1% of SkM tissue-derived EVs from either WT or mT mice were mG positive. By contrast, SkM tissue-derived EVs from SkM-mG/mT mice consistently displayed eGFP expression (30.54 ± 18.4% of events). An overlay of representative experiments from WT and SkM-mG/mT mice is shown in **Figure 5C**. In addition to single positive mG events, SkM tissue-derived EVs from SkM-mG/mT mice also had on average 14.28 ± 6.2% dual positive (mG and mT) events but only 1.38 ± 0.4% single positive (mT only) events.

To confirm that mG-containing EVs are specifically secreted by SkM myofibers, we also measured tdTomato and eGFP in WAT-derived EVs. mG was not detected in EVs secreted by WAT of SkM-mG/mT mice (**Figure 5D**). Finally, detergent lysis of EVs from SkM-mG/mT mice with 0.1% NP-40 eliminated fluorescence thus confirming the presence of endogenous fluorescent proteins in EVs secreted *ex vivo* (**Figure 5E**). Collectively, these data suggest that SkM myofibers account for a significant proportion of SkM tissue-derived EVs secreted *ex vivo*.

### Evidence for SkM-derived EVs in plasma and non-SkM cells in vivo

With evidence that mT and SkM myofiber-derived mG can be incorporated into EVs secreted *ex vivo*, we next wanted to examine to what extent SkM myofiber-derived EVs reach blood plasma *in vivo*. To address this question, we isolated EVs from plasma of WT, mT or SkM-mG/mT mice using size exclusion chromatography, Exoquick plasma kit or Exoquick-TC (to match our *ex vivo* isolation conditions). Similar results were obtained using all three methods; the results shown here are representative experiments from plasma EVs isolated using Exoquick-TC. In WT plasma (**Figure 6A**), less than 1% of EVs were positive for mT or mG. In mT plasma (**Figure 6B**), 6.0% of EVs were mT positive but no mG positive events were detected. Finally, plasma EVs from SkM-mG/mT mice contained 4.6% mT positive events and 0.26% mG positive events (**Figure 6C**).

Since the abundance of mG^+^ EVs was quite low in plasma, we isolated splenocytes, whole liver cells and erythrocytes to determine the extent to which SkM myofiber-derived mG might incorporate into distant tissues. As shown in **Figure 6D**, splenocytes robustly express mT as expected. Furthermore, mG does appear to be present, albeit at very low levels in splenocytes (**Figure 6E**). We confirmed the same robust presence of mT (**Figure 6F**) but modest presence of mG (**Figure 6G**) in whole liver cell preparations. Finally, erythrocytes also expressed mT (**Figure 6H**) but not mG (**Figure 6I**). These data suggest that SkM myofiber-derived mG may reach the plasma and distal tissues like the spleen and liver but are not taken up by circulating erythrocytes. Collectively, these data indicate that the amount of mG found in plasma is far less than that found in conditioned media from SkM explants, suggesting that SkM myofibers may not be a major contributor to EVs in circulation.

## Discussion

Defining the origin, abundance and function of EVs from specific cells and tissues is vital for using EVs as biomarkers and engineered therapies. There are a number of factors that dictate how much a given cell type will contribute to circulating EV abundance, including: tissue mass, tissue EV secretion capacity, EV access to circulation and EV clearance. None of these factors have been well defined, emphasizing the need for targeted approaches and novel methodologies. SkM and WAT are, by mass, two of the largest tissues in the body and would therefore be predicted to be major contributors to circulating EV abundance. Using both curated human data and an *ex vivo* tissue culture assay we demonstrate that, on a per mass basis, SkM tissue secretes more EVs than WAT. We show that intrinsic metabolic capacity is a determinant of SkM EV secretion capacity that may explain differences in EV secretion between SkM and WAT. Using spectral flow cytometry, we demonstrate that SkM tissue-derived EVs secreted *ex vivo* originate, at least in part, from SkM myofibers. Despite the robust ability of SkM myofibers to secrete EVs *ex vivo*, few (if any) SkM myofiber-derived EVs were found in blood plasma or distal organs. Therefore, despite its large mass and EV secretion capacity, our data do not support SkM myofibers being a major source of circulating EVs in healthy sedentary mice. Instead, our findings suggest the possibility that EVs derived from SkM myofibers and possibly other non-vascular cells exist in other extracellular compartments like the interstitial space.

Much of the data described in this report was obtained with an *ex vivo* EV secretion assay using mouse tissue explants. There is precedent for the use of similar approaches in both SkM (1, 25) and WAT (37). For the studies described here, there were multiple advantages to using this *ex vivo* approach. First, this method allowed us to directly compare EV secretion between tissues of individual mice. The within-subject design used here eliminated the contribution of variance between animals and permitted the use of pair-wise statistical comparisons. Second, the quality of the data presented here was strengthened by normalizing EV secretion to tissue mass, which allowed us to directly compare EV secretion capacity while eliminating the variance associated with differences in tissue input. Third, this method allowed for the study of EV secretion from cells existing in their native extracellular environment. This is important because EVs are known to respond to and interact with the extracellular matrix (13), which serves not only as a scaffold but also a communication interface that is inextricably linked to cell signaling and metabolism (19). Finally, by studying SkM tissue instead of cell monocultures, the contribution of SkM myofibers to EV secretion relative to other tissue resident cells could be assessed. This is important since resident SkM progenitor cells and fibroblasts are also known to secrete EVs (10, 23). While we contend that our tissue-based approach was the best available model for the questions addressed here, the limitations should also be noted. For example, dissection of SkM tissue damages myofibers and increases the risk of cellular contamination. Indeed, we did detect GM130, a Golgi-associated protein, in both SkM and WAT-derived EVs (**Figure 1B and C**). It is unclear to what extent cellular contamination may affect our results, but mean (**Figures 1H, 2B and 2E**) and modal (**Figures 1I, 2C and 2F**) EV sizes determined with NTA in four different datasets demonstrate that the vast majority of particles in our samples are “small” EVs (< 200 nm diameter) as defined by the International Society for Extracellular Vesicles (ISEV) (31). While it is apparent that tissue explant-derived EVs present challenges with regard to EV purity, the methods described here represent an important advance in this underdeveloped area of research.

We found using publicly available human tissue and cell data that human SkM and myofibers contain more of the molecular machinery needed to synthesize EVs than WAT and white adipocytes (**Table 1**). Consistent with this observation, we found that EV secretion capacity is greater in the mixed vastus medialis SkM compared to epididymal WAT *ex vivo* (**Figure 1G**). There are a few distinguishing features between SkM and WAT that could explain the strikingly greater rate of EV secretion in SkM compared to WAT. We examined two of these possibilities in this report. First, we demonstrated that spontaneous SkM contraction, which we inhibited with blebbistatin, did not explain greater EV secretion in SkM compared to WAT (**Figure 2A**). The blebbistatin concentration used in this study (10 μM) was chosen because we have shown it to be sufficient to prevent contraction in mouse and human SkM explants in previous studies (27). The lack of a decrease in EV abundance is intriguing in part because, at higher concentrations, blebbistatin has been shown to inhibit EV secretion from tumor cells (at 50 μM) (2) and inhibit the release of small (~30 nm) cholesterol-rich particles from macrophages (at 30 μM) (14). These previous studies suggest that blebbistatin might decrease SkM EV secretion at higher doses, but since this was observed in non-contracting cells, the effect would likely be independent of SkM contraction. Another possibility to explain these tissue-specific differences in EV secretion between SkM and WAT is intrinsic metabolic capacity. SkM is a far more metabolically active tissue compared to WAT so we hypothesized that tissues with greater intrinsic metabolic capacity would have a greater capacity for EV secretion. Previous work indicated that oxidative SkM secretes more EV protein than glycolytic SkM (25), however, this study determined EV quantity based on acetylcholinesterase which may not be a suitable index of EV abundance (20). Here, we confirmed this hypothesis (and those previous findings) using NTA to directly compare EV secretion between soleus, a highly oxidative muscle, and plantaris, which is more glycolytic (**Figure 2D**). These data collectively demonstrate that SkM has a higher capacity to secrete EVs than WAT and that intrinsic metabolic capacity predicts SkM EV secretion capacity *ex vivo* and may explain differences in EV secretion between SkM and WAT.

We observed an abundance of fluorescently-labeled EVs (mT and/or mG) in both SkM and WAT *ex vivo* (**Figure 5**) but far fewer (if any) in the blood plasma of the same mice (**Figure 6**). To our knowledge, we are the first group to examine the abundance of SkM myofiber-derived EVs using a transgenic mouse model. However, there are at least two previous reports that used fluorescent reporter mice to examine the abundance of other non-vascular cell-derived EVs in circulation. Pua et al. (28) generated mG/mT mice where eGFP was expressed specifically in pulmonary epithelial cells. Using differential ultracentrifugation, the authors detected an abundance of mT and mG-labeled EVs in brochoalveolar lavage fluid but fluorescent EVs were absent in serum. The findings described here (plasma EVs isolated using precipitation or SEC), by Pua et al. (serum EVs isolated using differential ultracentrifugation) (28) and Flaherty et al. (ultrafiltration and size exclusion chromatography (7) support two complementary and emerging paradigms. First, the results of these studies are consistent with previous work showing that EVs represent only a small fraction of EV-sized particles found in plasma (12, 38). Since cell-specific fluorescent EVs can be detected in abundance *ex vivo*, our results suggest that EVs secreted by non-vascular cells like SkM myofibers may be largely restricted to specific extracellular compartments. The recent discovery and characterization of matrix-bound nanovesicles in pig urinary bladder and dermis (15, 16) suggest that other tissues (i.e. SkM and WAT) may also have EVs embedded in the extracellular matrix. Future studies might utilize the SkM-mG/mT mouse or other endogenous fluorescent reporter mouse models to determine the extent to which SkM myofiber-derived EVs secreted *in vivo* exist in distinct extracellular compartments.

The lack of fluorescent EVs in blood plasma (**Figure 6**) may be explained primarily by co-isolated lipoproteins and/or protein aggregates (12, 38), but there are at least two other possibilities to be considered. First, EVs secreted from SkM myofibers may be modified as they travel through the interstitial space across the vascular endothelium. While there are reports of EVs being taken up by endothelial cells (36), to our knowledge, there are no reports of basal-to-apical flux of EVs through endothelial cells or the gap junctions that perforate the endothelium. Even small EVs like exosomes (50-200 nm diameter) are much larger than gap junctions (~ 2 nm diameter) and fairly rigid, suggesting that their transport is more likely to occur through endothelial cells rather than via gap junctions. Second, monocytes like macrophages may selectively extract and/or modify EVs in the circulation. There is some evidence for monocytic handling of EVs in blood (17) that is consistent with their role in detoxification. Monocytes also exist in most other tissue compartments (including SkM and WAT) but do not appear to preclude the detection of fluorescent EVs in our *ex vivo* experiments or those by Flaherty et al. (7). To what extent EVs may be modified traveling through the endothelium or following uptake by monocytes is unknown but could be addressed in future studies. The continued development and application of fluorescent reporter mice like the mG/mT mouse and others (i.e. conditional CD63-GFP mouse (22)) will further elucidate the distribution of cell-specific EVs *in vivo*.

An improved understanding of EV secretion by different tissues and their biodistribution *in vivo* is needed to realize the full potential of EVs as biomarkers and therapeutics. Our findings contribute to this broad objective by: 1) confirming that SkM myofibers secrete EVs that can be detected *ex vivo* and possibly *in vivo*, 2) revealing that metabolic activity is a determinant of EV secretion both within and between tissues and 3) that the rate of tissue EV secretion *ex vivo* does not necessarily reflect circulating EV abundance *in vivo*.

## Acknowledgements

The authors wish to thank the following sources. Colorado State University Flow Cytometry and Cell Sorting Facility for their help in method development, optimization and sample analysis. American Heart Association for extramural support through an Innovative Project Award (IPA1834110052 to DSL).

## References

1. Aswad H, Forterre A, Wiklander OP, Vial G, Danty-Berger E, Jalabert A, Lamaziere A, Meugnier E, Pesenti S, Ott C, Chikh K, El-Andaloussi S, Vidal H, Lefai E, Rieusset J, and Rome S. Exosomes participate in the alteration of muscle homeostasis during lipid-induced insulin resistance in mice. Diabetologia 57: 2155–2164, 2014.

2. Brassart B, Da Silva J, Donet M, Seurat E, Hague F, Terryn C, Velard F, Michel J, Ouadid-Ahidouch H, Monboisse JC, Hinek A, Maquart FX, Ramont L, and Brassart-Pasco S. Tumour cell blebbing and extracellular vesicle shedding: key role of matrikines and ribosomal protein SA. Br J Cancer 120: 453–465, 2019.

3. Connolly KD, Wadey RM, Mathew D, Johnson E, Rees DA, and James PE. Evidence for Adipocyte-Derived Extracellular Vesicles in the Human Circulation. Endocrinology 159: 3259–3267, 2018.

4. De Gasperi R, Hamidi S, Harlow LM, Ksiezak-Reding H, Bauman WA, and Cardozo CP. Denervation-related alterations and biological activity of miRNAs contained in exosomes released by skeletal muscle fibers. Sci Rep 7: 12888, 2017.

5. Deng ZB, Poliakov A, Hardy RW, Clements R, Liu C, Liu Y, Wang J, Xiang X, Zhang S. Zhuang X, Shah SV, Sun D, Michalek S, Grizzle WE, Garvey T, Mobley J, and Zhang HG. Adipose tissue exosome-like vesicles mediate activation of macrophage-induced insulin resistance. Diabetes 58: 2498–2505, 2009.

6. Diaz G, Bridges C, Lucas M, Cheng Y, Schorey JS, Dobos KM, and Kruh-Garcia NA. Protein Digestion, Ultrafiltration, and Size Exclusion Chromatography to Optimize the Isolation of Exosomes from Human Blood Plasma and Serum. J Vis Exp 2018.

7. Flaherty SE, 3rd, Grijalva A, Xu X, Ables E, Nomani A, and Ferrante AW, Jr. A lipase-independent pathway of lipid release and immune modulation by adipocytes. Science 363: 989–993, 2019.

8. Forterre A, Jalabert A, Berger E, Baudet M, Chikh K, Errazuriz E, De Larichaudy J, Chanon S, Weiss-Gayet M, Hesse AM, Record M, Geloen A, Lefai E, Vidal H, Couté Y, and Rome S. Proteomic analysis of C2C12 myoblast and myotube exosome-like vesicles: a new paradigm for myoblast-myotube cross talk? PLoS One 9: e84153, 2014.

9. Frühbeis C, Helmig S, Tug S, Simon P, and Krämer-Albers E-M. Physical exercise induces rapid release of small extracellular vesicles into the circulation. Journal of extracellular vesicles 4: 28239–28239, 2015.

10. Fry CS, Kirby TJ, Kosmac K, McCarthy JJ, and Peterson CA. Myogenic Progenitor Cells Control Extracellular Matrix Production by Fibroblasts during Skeletal Muscle Hypertrophy. Cell Stem Cell 20: 56–69, 2017.

11. Groot Kormelink T, Arkesteijn GJ, Nauwelaers FA, van den Engh G, Nolte-’t Hoen EN, and Wauben MH. Prerequisites for the analysis and sorting of extracellular vesicle subpopulations by high-resolution flow cytometry. Cytometry A 89: 135–147, 2016.

12. György B, Módos K, Pállinger E, Pálóczi K, Pásztói M, Misják P, Deli MA, Sipos A, Szalai A, Voszka I, Polgár A, Tóth K, Csete M, Nagy G, Gay S, Falus A, Kittel A, and Buzás EI. Detection and isolation of cell-derived microparticles are compromised by protein complexes resulting from shared biophysical parameters. Blood 117: e39–48, 2011.

13. Hoshino A, Costa-Silva B, Shen TL, Rodrigues G, Hashimoto A, Tesic Mark M, Molina H, Kohsaka S, Di Giannatale A, Ceder S, Singh S, Williams C, Soplop N, Uryu K, Pharmer L, King T, Bojmar L, Davies AE, Ararso Y, Zhang T, Zhang H, Hernandez J, Weiss JM, Dumont-Cole VD, Kramer K, Wexler LH, Narendran A, Schwartz GK, Healey JH, Sandstrom P, Labori KJ, Kure EH, Grandgenett PM, Hollingsworth MA, de Sousa M, Kaur S, Jain M, Mallya K, Batra SK, Jarnagin WR, Brady MS, Fodstad O, Muller V, Pantel K, Minn AJ, Bissell MJ, Garcia BA, Kang Y, Rajasekhar VK, Ghajar CM, Matei I, Peinado H, Bromberg J, and Lyden D. Tumour exosome integrins determine organotropic metastasis. Nature 527: 329–335, 2015.

14. Hu X, Weston TA, He C, Jung RS, Heizer PJ, Young BD, Tu Y, Tontonoz P, Wohlschlegel JA, Jiang H, Young SG, and Fong LG. Release of cholesterol-rich particles from the macrophage plasma membrane during movement of filopodia and lamellipodia. Elife 8: 2019.

15. Huleihel L, Hussey GS, Naranjo JD, Zhang L, Dziki JL, Turner NJ, Stolz DB, and Badylak SF. Matrix-bound nanovesicles within ECM bioscaffolds. Sci Adv 2: e1600502, 2016.

16. Hussey GS, Pineda Molina C, Cramer MC, Tyurina YY, Tyurin VA, Lee YC, El-Mossier SO, Murdock MH, Timashev PS, Kagan VE, and Badylak SF. Lipidomics and RNA sequencing reveal a novel subpopulation of nanovesicle within extracellular matrix biomaterials. Sci Adv 6: eaay4361, 2020.

17. Imai T, Takahashi Y, Nishikawa M, Kato K, Morishita M, Yamashita T, Matsumoto A, Charoenviriyakul C, and Takakura Y. Macrophage-dependent clearance of systemically administered B16BL6-derived exosomes from the blood circulation in mice. J Extracell Vesicles 4: 26238, 2015.

18. Jalabert A, Vial G, Guay C, Wiklander OP, Nordin JZ, Aswad H, Forterre A, Meugnier E, Pesenti S, Regazzi R, Danty-Berger E, Ducreux S, Vidal H, El-Andaloussi S, Rieusset J, and Rome S. Exosome-like vesicles released from lipid-induced insulin-resistant muscles modulate gene expression and proliferation of beta recipient cells in mice. Diabetologia 59: 1049–1058, 2016.

19. Lark DS, and Wasserman DH. Meta-fibrosis links positive energy balance and mitochondrial metabolism to insulin resistance. F1000Research 6: 1758, 2017.

20. Liao Z, Jaular LM, Soueidi E, Jouve M, Muth DC, Schøyen TH, Seale T, Haughey NJ, Ostrowski M, Théry C, and Witwer KW. Acetylcholinesterase is not a generic marker of extracellular vesicles. J Extracell Vesicles 8: 1628592, 2019.

21. McVey MJ, Spring CM, and Kuebler WM. Improved resolution in extracellular vesicle populations using 405 instead of 488 nm side scatter. J Extracell Vesicles 7: 1454776, 2018.

22. Men Y, Yelick J, Jin S, Tian Y, Chiang MSR, Higashimori H, Brown E, Jarvis R, and Yang Y. Exosome reporter mice reveal the involvement of exosomes in mediating neuron to astroglia communication in the CNS. Nature Communications 10: 4136, 2019.

23. Murach KA, Vechetti IJ, Jr., Van Pelt DW, Crow SE, Dungan CM, Figueiredo VC, Kosmac K, Fu X, Richards CI, Fry CS, McCarthy JJ, and Peterson CA. Fusion-Independent Satellite Cell Communication to Muscle Fibers During Load-Induced Hypertrophy. Function (Oxf) 1: zqaa009, 2020.

24. Muzumdar MD, Tasic B, Miyamichi K, Li L, and Luo L. A global double-fluorescent Cre reporter mouse. Genesis 45: 593–605, 2007.

25. Nie Y, Sato Y, Garner RT, Kargl C, Wang C, Kuang S, Gilpin CJ, and Gavin TP. Skeletal muscle-derived exosomes regulate endothelial cell functions via reactive oxygen species-activated nuclear factor-κB signalling. Exp Physiol 104: 1262–1273, 2019.

26. Noren Hooten N, and Evans MK. Extracellular vesicles as signaling mediators in type 2 diabetes mellitus. Am J Physiol Cell Physiol 318: C1189–c1199, 2020.

27. Perry CG, Kane DA, Lin CT, Kozy R, Cathey BL, Lark DS, Kane CL, Brophy PM, Gavin TP, Anderson EJ, and Neufer PD. Inhibiting myosin-ATPase reveals a dynamic range of mitochondrial respiratory control in skeletal muscle. Biochem J 437: 215–222, 2011.

28. Pua HH, Happ HC, Gray CJ, Mar DJ, Chiou NT, Hesse LE, and Ansel KM. Increased Hematopoietic Extracellular RNAs and Vesicles in the Lung during Allergic Airway Responses. Cell Rep 26: 933–944.e934, 2019.

29. Racine ML, and Dinenno FA. Reduced deformability contributes to impaired deoxygenation-induced ATP release from red blood cells of older adult humans. J Physiol 597: 4503–4519, 2019.

30. Rome S, Forterre A, Mizgier ML, and Bouzakri K. Skeletal Muscle-Released Extracellular Vesicles: State of the Art. Front Physiol 10: 929, 2019.

31. Théry C, Witwer KW, Aikawa E, Alcaraz MJ, Anderson JD, Andriantsitohaina R, Antoniou A, Arab T, Archer F, Atkin-Smith GK, Ayre DC, Bach J-M, Bachurski D, Baharvand H, Balaj L, Baldacchino S, Bauer NN, Baxter AA, Bebawy M, Beckham C, Bedina Zavec A, Benmoussa A, Berardi AC, Bergese P, Bielska E, Blenkiron C, Bobis-Wozowicz S, Boilard E, Boireau W, Bongiovanni A, Borràs FE, Bosch S, Boulanger CM, Breakefield X, Breglio AM, Brennan MÁ, Brigstock DR, Brisson A, Broekman MLD, Bromberg JF, Bryl-Górecka P, Buch S, Buck AH, Burger D, Busatto S, Buschmann D, Bussolati B, Buzás EI, Byrd JB, Camussi G, Carter DRF, Caruso S, Chamley LW, Chang Y-T, Chen C, Chen S, Cheng L, Chin AR, Clayton A, Clerici SP, Cocks A, Cocucci E, Coffey RJ, Cordeiro-da-Silva A, Couch Y, Coumans FAW, Coyle B, Crescitelli R, Criado MF, D’Souza-Schorey C, Das S, Datta Chaudhuri A, de Candia P, De Santana EF, De Wever O, del Portillo HA, Demaret T, Deville S, Devitt A, Dhondt B, Di Vizio D, Dieterich LC, Dolo V, Dominguez Rubio AP, Dominici M, Dourado MR, Driedonks TAP, Duarte FV, Duncan HM, Eichenberger RM, Ekström K, El Andaloussi S, Elie-Caille C, Erdbrügger U, Falcón-Pérez JM, Fatima F, Fish JE, Flores-Bellver M, Försönits A, Frelet-Barrand A, Fricke F, Fuhrmann G, Gabrielsson S, Gámez-Valero A, Gardiner C, Gärtner K, Gaudin R, Gho YS, Giebel B, Gilbert C, Gimona M, Giusti I, Goberdhan DCI, Görgens A, Gorski SM, Greening DW, Gross JC, Gualerzi A, Gupta GN, Gustafson D, Handberg A, Haraszti RA, Harrison P, Hegyesi H, Hendrix A, Hill AF, Hochberg FH, Hoffmann KF, Holder B, Holthofer H, Hosseinkhani B, Hu G, Huang Y, Huber V, Hunt S, Ibrahim AG-E, Ikezu T, Inal JM, Isin M, Ivanova A, Jackson HK, Jacobsen S, Jay SM, Jayachandran M, Jenster G, Jiang L, Johnson SM, Jones JC, Jong A, Jovanovic-Talisman T, Jung S, Kalluri R, Kano S-i, Kaur S, Kawamura Y, Keller ET, Khamari D, Khomyakova E, Khvorova A, Kierulf P, Kim KP, Kislinger T, Klingeborn M, Klinke DJ, Kornek M, Kosanović MM, Kovács ÁF, Krämer-Albers E-M, Krasemann S, Krause M, Kurochkin IV, Kusuma GD, Kuypers S, Laitinen S, Langevin SM, Languino LR, Lannigan J, Lässer C, Laurent LC, Lavieu G, Lázaro-Ibáñez E, Le Lay S, Lee M-S, Lee YXF, Lemos DS, Lenassi M, Leszczynska A, Li ITS, Liao K, Libregts SF, Ligeti E, Lim R, Lim SK, Linē A, Linnemannstöns K, Llorente A, Lombard CA, Lorenowicz MJ, Lörincz ÁM, Lötvall J, Lovett J, Lowry MC, Loyer X, Lu Q, Lukomska B, Lunavat TR, Maas SLN, Malhi H, Marcilla A, Mariani J, Mariscal J, Martens-Uzunova ES, Martin-Jaular L, Martinez MC, Martins VR, Mathieu M, Mathivanan S, Maugeri M, McGinnis LK, McVey MJ, Meckes DG, Meehan KL, Mertens I, Minciacchi VR, Möller A, Møller Jørgensen M, Morales-Kastresana A, Morhayim J, Mullier F, Muraca M, Musante L, Mussack V, Muth DC, Myburgh KH, Najrana T, Nawaz M, Nazarenko I, Nejsum P, Neri C, Neri T, Nieuwland R, Nimrichter L, Nolan JP, Nolte-’t Hoen ENM, Noren Hooten N, O’Driscoll L, O’Grady T, O’Loghlen A, Ochiya T, Olivier M, Ortiz A, Ortiz LA, Osteikoetxea X, Østergaard O, Ostrowski M, Park J, Pegtel DM, Peinado H, Perut F, Pfaffl MW, Phinney DG, Pieters BCH, Pink RC, Pisetsky DS, Pogge von Strandmann E, Polakovicova I, Poon IKH, Powell BH, Prada I, Pulliam L, Quesenberry P, Radeghieri A, Raffai RL, Raimondo S, Rak J, Ramirez MI, Raposo G, Rayyan MS, Regev-Rudzki N, Ricklefs FL, Robbins PD, Roberts DD, Rodrigues SC, Rohde E, Rome S, Rouschop KMA, Rughetti A, Russell AE, Saá P, Sahoo S, Salas-Huenuleo E, Sánchez C, Saugstad JA, Saul MJ, Schiffelers RM, Schneider R, Schøyen TH, Scott A, Shahaj E, Sharma S, Shatnyeva O, Shekari F, Shelke GV, Shetty AK, Shiba K, Siljander PRM, Silva AM, Skowronek A, Snyder OL, Soares RP, Sódar BW, Soekmadji C, Sotillo J, Stahl PD, Stoorvogel W, Stott SL, Strasser EF, Swift S, Tahara H, Tewari M, Timms K, Tiwari S, Tixeira R, Tkach M, Toh WS, Tomasini R, Torrecilhas AC, Tosar JP, Toxavidis V, Urbanelli L, Vader P, van Balkom BWM, van der Grein SG, Van Deun J, van Herwijnen MJC, Van Keuren-Jensen K, van Niel G, van Royen ME, van Wijnen AJ, Vasconcelos MH, Vechetti IJ, Veit TD, Vella LJ, Velot É, Verweij FJ, Vestad B, Viñas JL, Visnovitz T, Vukman KV, Wahlgren J, Watson DC, Wauben MHM, Weaver A, Webber JP, Weber V, Wehman AM, Weiss DJ, Welsh JA, Wendt S, Wheelock AM, Wiener Z, Witte L, Wolfram J, Xagorari A, Xander P, Xu J, Yan X, Yáñez-Mó M, Yin H, Yuana Y, Zappulli V, Zarubova J, Žėkas V, Zhang J-y, Zhao Z, Zheng L, Zheutlin AR, Zickler AM, Zimmermann P, Zivkovic AM, Zocco D, and Zuba-Surma EK. Minimal information for studies of extracellular vesicles 2018 (MISEV2018): a position statement of the International Society for Extracellular Vesicles and update of the MISEV2014 guidelines. Journal of Extracellular Vesicles 7: 1535750, 2018.

32. Thomou T, Mori MA, Dreyfuss JM, Konishi M, Sakaguchi M, Wolfrum C, Rao TN, Winnay JN, Garcia-Martin R, Grinspoon SK, Gorden P, and Kahn CR. Adipose-derived circulating miRNAs regulate gene expression in other tissues. Nature 542: 450–455, 2017.

33. van der Pol E, van Gemert MJ, Sturk A, Nieuwland R, and van Leeuwen TG. Single vs. swarm detection of microparticles and exosomes by flow cytometry. J Thromb Haemost 10: 919–930, 2012.

34. Welsh JA, Van Der Pol E, Arkesteijn GJA, Bremer M, Brisson A, Coumans F, Dignat-George F, Duggan E, Ghiran I, Giebel B, Görgens A, Hendrix A, Lacroix R, Lannigan J, Libregts SFWM, Lozano-Andrés E, Morales-Kastresana A, Robert S, De Rond L, Tertel T, Tigges J, De Wever O, Yan X, Nieuwland R, Wauben MHM, Nolan JP, and Jones JC. MIFlowCyt-EV: a framework for standardized reporting of extracellular vesicle flow cytometry experiments. Journal of Extracellular Vesicles 9: 1713526, 2020.

35. Whitham M, Parker BL, Friedrichsen M, Hingst JR, Hjorth M, Hughes WE, Egan CL, Cron L, Watt KI, Kuchel RP, Jayasooriah N, Estevez E, Petzold T, Suter CM, Gregorevic P, Kiens B, Richter EA, James DE, Wojtaszewski JFP, and Febbraio MA. Extracellular Vesicles Provide a Means for Tissue Crosstalk during Exercise. Cell Metab 27: 237–251.e234, 2018.

36. Wu SF, Noren Hooten N, Freeman DW, Mode NA, Zonderman AB, and Evans MK. Extracellular vesicles in diabetes mellitus induce alterations in endothelial cell morphology and migration. J Transl Med 18: 230, 2020.

37. Xie Z, Wang X, Liu X, Du H, Sun C, Shao X, Tian J, Gu X, Wang H, Tian J, and Yu B. Adipose-Derived Exosomes Exert Proatherogenic Effects by Regulating Macrophage Foam Cell Formation and Polarization. J Am Heart Assoc 7: 2018.

38. Zhang X, Borg EGF, Liaci AM, Vos HR, and Stoorvogel W. A novel three step protocol to isolate extracellular vesicles from plasma or cell culture medium with both high yield and purity. Journal of Extracellular Vesicles 9: 1791450, 2020.

